# MechanoMR microparticle (M^3^) sensors reveal dynamic stress loading as a driver of epithelial-mesenchymal transition

**DOI:** 10.64898/2025.12.10.693598

**Authors:** Minji An, Ri Yu, Annie Lin, Jinho Jang, Jiyeon Ohk, Young-Joo Kim, Hosung Jung, Minsuk Kwak, Young-wook Jun, Jinwoo Cheon

## Abstract

The dynamic mechanical response of tissues underlies their physiological function, yet direct, quantitative measurement of tissue stress *in vivo* has remained a major challenge. Here, we introduce the <u>m</u>echano<u>M</u>R <u>m</u>icroparticle (M^3^, “M-cube”) sensor, a hybrid soft-matter/nanoparticle probe that integrates directly into tissue mechanical networks while transducing local stress into quantitative magnetic resonance (MR) readouts with single-particle resolution. We demonstrate the utility of this platform across diverse model systems, including tumor spheroids, Xenopus embryos, and mouse xenografts, where the M^3^ sensor enables noninvasive, spatiotemporally resolved mapping of tissue stress dynamics during cancer development. Using this approach, we reveal that epithelial-mesenchymal transition (EMT) is accompanied by distinctive stress-remodeling patterns observable *in vivo*. Strikingly, we find that abrupt stress increases, rather than cumulative or peak stress magnitude, are the key determinants of EMT induction in cancer cells within the tumor microenvironment. Transcriptomic profiling under controlled stress-loading dynamics shows that sustained yet gradual stress escalation activates cytoprotective antioxidation pathways (e.g., FOXO/AMPK) that reinforce epithelial stability, whereas acute stress surges overwhelm these defense mechanisms, predisposing cells toward mesenchymal reprogramming. These findings establish the M^3^ sensor as a broadly applicable technology for linking dynamic mechanical cues to cell-state transitions in development, homeostasis, and disease.

---

Tissues continuously generate and experience mechanical forces that are essential for their physiological function and, when dysregulated, drive disease^1,2^. Mechanical cues shape diverse processes in tissue development^3^, cardiovascular homeostasis^4^, and tumor progression, where accumulating internal stress modulates cell behavior, invasiveness, and metastatic potential^5^. Given their central importance, substantial efforts over the past decades have been devoted to developing methods to quantify tissue mechanics. These include contact probes^6^, optical or magnetic tweezers^7^, laser ablation^8,9^, traction-force microscopy^10^, molecular tension sensors^11–13^, stress-relaxation mapping^14^, interferometric approaches such as optical coherence tomography^15^, elastography^16^, and soft-matter inclusion methods such as hydrogel beads and droplets^17–21^. These technologies have transformed *in vitro* mechanobiology and, in some cases, entered clinical use^22^. Yet *in vivo* tissue mechanics remain challenging: existing approaches are limited by restricted applicability to transparent tissues^18,23^, invasiveness and single-point readouts^14,21^, or an inability to probe the actual stress state rather than bulk materials properties such as stiffness^16,24^.

To overcome these barriers, we sought to develop a method that converts local mechanical stress within tissues *in vivo* into quantitative readouts with high spatiotemporal resolution and precision, which can be readily sensed using established imaging modalities. Here, we devise such a system, the mechanoMR microparticle (M^3^, “M-cube”) sensor, which integrates a deformable alginate hydrogel with embedded magnetic nanoparticles to transduce local stress into single-particle MR signals. Using this platform, we investigate spatiotemporal dynamics of mechanical stress during tumor development and differentiation in diverse *in vitro* and *in vivo* model systems. We also correlate local stress distributions with cellular responses within their native mechanical microenvironments. Lastly, through transcriptomics analysis under defined mechanical stress conditions, we examine how dynamic tissue stress landscapes perturb molecular programs of cancer cells and prime them for EMT.

## M^3^ sensor: synthesis, design, and working principle

The M^3^ sensor, consisting of 70 µm alginate microparticles (0.6%) embedded with uniformly distributed zinc-doped ferrite nanoparticles (Zn_0.4_Fe_2.6_O_4_, 13 nm, was synthesized by microfluidic in-droplet click chemistry (**Fig. 1a**, details in Methods). This sensor is engineered on the basis of two key technologies. 1) hydrogel inclusion stress-mapping methods and 2) enhanced MR contrast effects by Zn_0.4_Fe_2.6_O_4_ nanoparticles^25^, enabling single microparticle MR imaging. The inclusion strategy directly integrates the microparticle into the 3D tissue stress tensor field, thereby enabling direct measurement of local solid stress by monitoring particle deformation (**Fig. 1b**), as exemplified in vertebrate body axis elongation^26,27^, spheroid stress mapping^18–20^, and *ex vivo* tumor stress mapping^14^. The M^3^ sensor fully exploits this unique capacity of the inclusion method for *in vivo* applications, previously restricted largely to optically transparent systems, through a simple and robust mechanism – mechanoMR transduction.

**Fig. 1.**
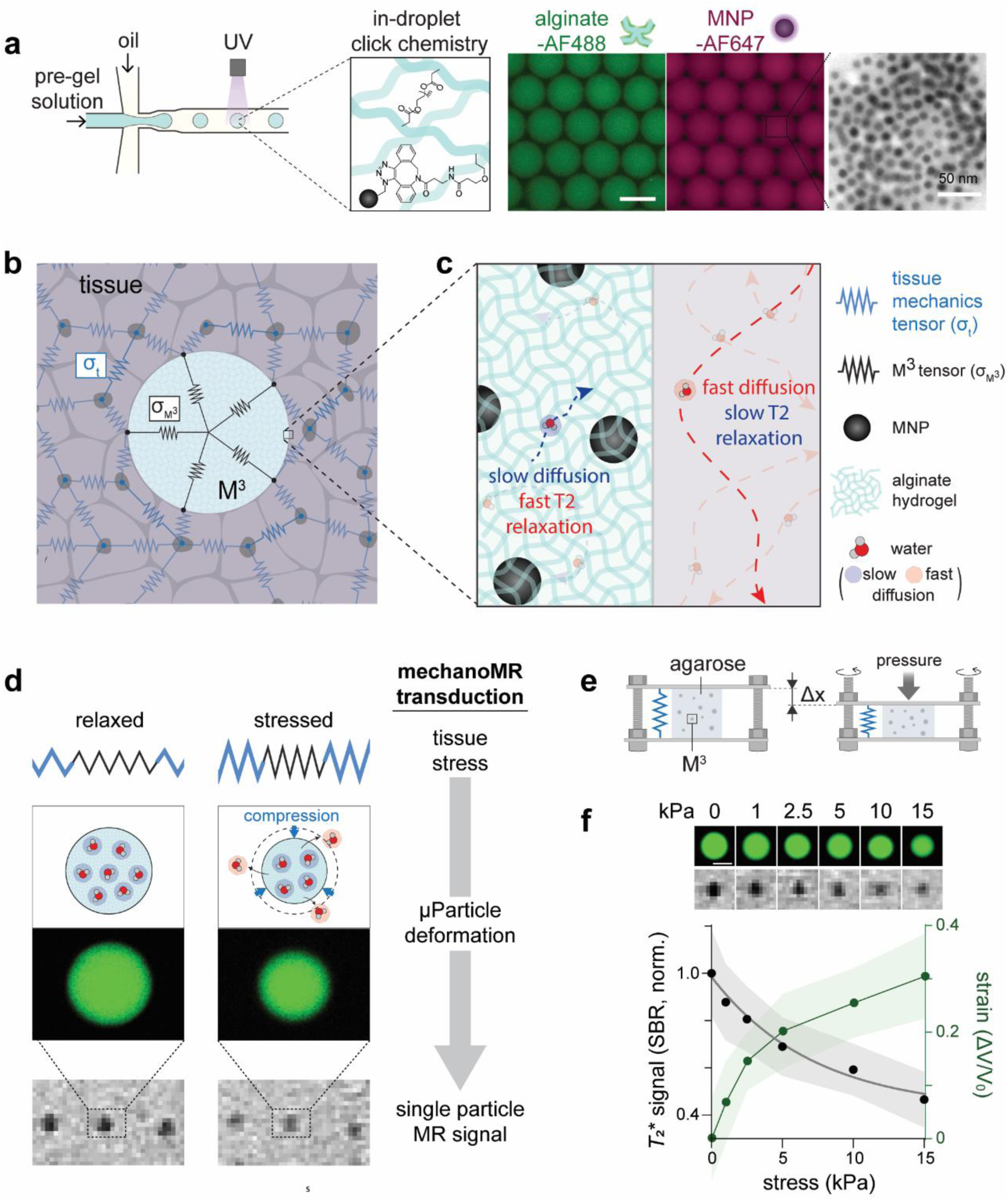
Synthesis, design, and working principle of the M^3^ sensor. **a,** Schematic of the microfluidic synthesis for M^3^ sensors using in-droplet click chemistry (left). Characterization of M^3^ sensors using fluorescence and transmission electron microscopies (TEM) (right). Scale bars, 50 μm and 50 nm (TEM). **b-d,** Schematics depicting M^3^ sensors mechanically integrated within tissue (**b**). The M^3^ sensor slows down water molecule diffusion within it, leading contrasted single-particle MR signals (**c**). When compressed due to tissue stress, single-particle MR signals of M^3^ sensors decrease via mechanoMR transduction (**d**). **e,** Setup for stress-MR signal calibration using a sandwich stress-application device. M^3^ sensors are dispersed in a cylindrical agarose gel block. Stress was generated by controlled vertical displacement (Δx) of the top plate. **f**, Representative confocal fluorescence microscopy (CFM) and MR images of M^3^ sensor under applied stress ranging from 0 to 15 kPa (top). Normalized *T*_2_* signal (left) and strain (right) profile of M^3^ sensors as a function of applied stress 0 to 15 kPa. (bottom) (*n* = 30 - 38). Data are presented as mean ± s.d.

The M^3^ sensor creates a dual environment for water molecules: slow diffusion due to hydrogel-backbone interactions, and fast spin-spin relaxation (*i.e.*, low *T*_2_, high *R*_2_) induced by embedded magnetic nanoparticles. Under local stress, the M^3^ sensor deforms, expels water molecules, and proportionally reduces *R*_2_ relaxivity via two coupled mechanisms: First, deformation decreases the fraction of slow-diffusing water molecules proximal to magnetic particles, thereby lowering *R*_2_^28^. Second, reduced interparticle spacing increases the likelihood that diffusing water protons encounter multiple nanoparticles during diffusion, which further lowers *R*_2_ via motional averaging (**Fig. 1c,d**). Together, these effects establish a robust inverse relationship between local mechanical stress and MR relaxivity (*R*_2_). Because MR imaging offers noninvasive, whole-tissue imaging, we hypothesize that dynamic tissue stress can be mapped through single-particle MR tracking of the M^3^ sensor.

## M^3^ sensor enables quantitative, single-particle resolution readout of environmental mechanical stress

To test this hypothesis, we implanted M^3^ sensors within a cylindrical agarose gel block (2%, Ø 5 mm × 10 mm height) and applied controlled compressive stress using a custom sandwich device (**Fig. 1e, Supplementary Fig. 1a,b**). The agarose gel block frequently serves as a tissue-mimetic medium that uniformly transmits stress throughout the gel under loading (**Supplementary Video 1**), and is compatible with both optical and MR imaging, enabling direct validation of sensor performance through deformation-MR signal comparison. To mimic physiologically relevant mechanical environment, we varied the applied stress between 0 and 15 kPa, and traced sensor volume changes and MR signals at the single particle level using confocal fluorescence microscopy (CFM) and *T*_2_*-weighted MR imaging, respectively. At no applied stress, we observed distinct dark spots in MRI corresponding individual microparticles, resolved at approximately 300 µm, indicating that M^3^ sensors generate strong (SN ratio: 9.5), reproducible (CV = 2.4%), and relatively homogeneous (s.d. = 10%) single-particle signals suitable for their intended applications (**Extended Data Fig. 1a,b**). These signals remained robust across environmental parameters – including pH, medium stiffness, medium type (agarose *vs*. matrigel), and up to four weeks of continuous monitoring – supporting their utility for long-term *in vivo* application (**Extended Data Fig. 1c-f**). Upon increasing applied stress, we detected a gradual decrease in single-particle MR signals that correlated with progressive particle compression. In contrast, PEGDA-based M^3^ sensors (E > 2.4 MPa), which are nearly incompressible within this stress range, exhibited no detectable signal change (**Extended Data Fig. 1g-i**). Repeated analysis of individual M^3^ sensors across varying applied stresses yielded a calibration curve defined by an empirical biexponential decay equation of *T*_2_* signal = 0.1344 e^−0.926S^ + 0.864 e^−0.0405S^, where S represents environmental stress (**Fig. 1f**). Together, these results validate the M^3^ sensor as a reliable platform for noninvasive, single-particle readout of mechanical stress, motivating its application in complex biological systems.

## Dynamic stress measurements during tumor spheroid growth and pathogenesis

We first applied the M^3^ sensor to quantify evolving mechanical stress in 3D tumor spheroid models, capturing how stress dynamics accompany tumor growth and pathogenesis. We chose 3D tumor spheroid models because they (1) recapitulate key mechanical features of the solid tumor microenvironment, including spatial gradients of proliferation, matrix resistance, and compressive buildup^29^ and (2) provide predictable *in vitro* platforms for interrogating tissue mechanical responses under controlled stimulation^30^. Additionally, when maintained below specific size thresholds, (3) spheroids permit *in situ* fluorescence detection of particle deformation alongside changes in cellular state. This feature enables correlated CFM-MRI measurements for rigorous sensor performance evaluation, while simultaneously allowing detailed investigation of tissue mechanics in relation to cellular state transitions. Hence, in all following experiments, we validated the results from M^3^ sensors by quantitatively tracking particle deformation in spheroids using this correlated imaging approach (**Extended Data Fig. 2a**).

We formed 2D arrays of M^3^-laden tumor spheroids by co-entrapping cancer cells and microparticles within microfabricated gelatin wells, followed by transferring the resulting cell-particle aggregates into a 3D matrigel matrix via sacrificial micromolding (**Fig. 2a,b, Supplementary Fig. 2**). This approach employs uniform microwell patterns with defined diameters and depths, thereby enabling synchronized spheroid growth and ensuring consistent matrix-imposed confinement, which standardizes the mechanical boundary conditions across samples. The resulting spheroid contained zero to two M^3^ sensors, with distinct dark MR signals detected exclusively in those harboring sensors (**Fig. 2b, Extended Data Fig. 2b**). For quantitative analysis of stress dynamics, we focused on tracking single-particle MR signals and, using the previously established calibration curve, converting these signals into local stress values for individual spheroids containing a single M^3^ sensor.

**Fig. 2.**
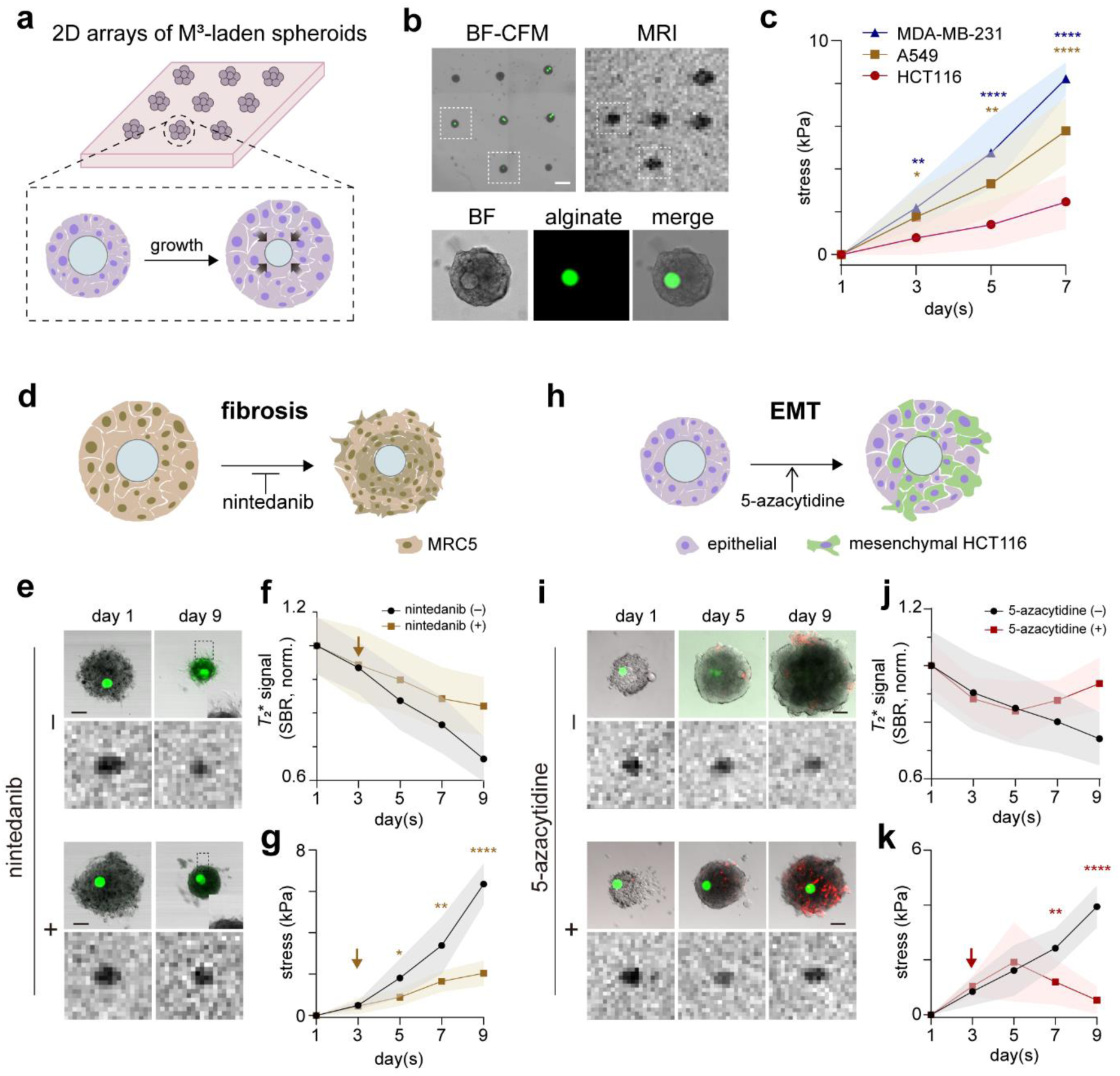
Longitudinal stress monitoring of developing tumor spheroids with M^3^ sensors. **a,** Schematic of 2D tumor spheroid arrays embedded with M^3^ sensors via sacrificial micromolding. **b,** The sensor integration into the spheroid array was detected by correlated CFM and MR imaging. (bottom; green, alginate). Scale bar, 500 µm. **c**, Quantitative stress mapping over the 7-day spheroid growth using the M^3^ sensor. Three cell lines (HCT116, A549, and MDA-MB-231) (*n* = 10, 13, and 11, respectively) were used for comparision. **d**-**g,** Stress development during fibrosis of MRC5 spheroids. A schematic (**d**), CFM, and MR images of fibrotic spheroids with or without fibrosis inhibitor, nintedanib (**e**). Scale bar, 100 μm. Normalized *T*_2_* signals (**f**) and stress (**g**) of the fibrotic spheroids with or without nintedanib treatment (*n* = 11 for (-) and *n* = 8 for (+)). **h**-**k,** Stress dynamics under epithelial-mesenchymal transition (EMT) of tumor spheroids. HCT116-vimentin-RFP reporter cells were used to detect EMT. A schematic (**h**), CFM, and MR images of HCT116 spheroids with or without EMT-inducer, 5-azacytidine (**i**). Scale bar, 100 μm. Normalized *T*_2_* signals (**j**) and stress (**k**) of HCT116 spheroids u with or without 5-azacytidine treatment (*n* = 9 for (-) and *n* = 12 for (+)). Data are presented as mean ± s.d. Statistical significance was determined by one-way ANOVA with Tukey’s multiple comparisons test for (**c**), and by two-tailed unpaired Student’s t-test for (**g**) and (**k**). ****P < 0.0001; **P < 0.01; *P < 0.05.

We examined three different types of tumor spheroids composed of colon (HCT116), lung (A549), and breast (MDA-MB-231) cancer cells. All three spheroid types exhibited a gradual decrease in single-particle MR signals during growth, suggesting progressive stress accumulation, but with distinct accumulation kinetics that reflect cell-type-specific mechanical remodeling (**Fig. 2c, Extended data Fig. 3**). We interpret these results as indicating that spheroids composed of stiffer cells (E_MDA-MB-231_ > E_A549_ > E_HCT116_) experienced more rapid compression buildup during proliferation within a confined matrix^31^. To further test this notion, we generated M^3^-laden spheroids composed of MRC5, a model cell line for pulmonary fibrosis, and monitored stress development over time (**Fig. 2d**). In contrast to the other spheroid types, which exhibited gradual size increases during growth, MRC5 spheroids progressively contracted, forming a more compact structure (**Fig. 2e**). Consistent with their strong contractile phenotype, MRC5 spheroids exhibited a markedly steeper trajectory of single particle MR signal reduction, *i.e.*, rapid stress accumulation, over time, suggesting that active matrix remodeling associated with fibrosis can accelerate compressive stress buildup (**Fig. 2e-g**). Inhibition of the fibrotic matrix remodeling with nintedanib treatment significantly slowed stress buildup, further confirming that active remodeling drives the elevated compressive stress dynamics observed in these spheroids (**Fig. 2e-g**). Finally, we applied the M^3^ sensors to investigate stress dynamics upon cell state transitions such as EMT. We formed M^3^-laden spheroids with vimentin-RFP reporter cells (HCT116-vimentin-RFP, **Supplementary Fig. 3**) and induced EMT by treating them with 5-azacytidine three days after spheroid seeding^32^ (**Fig. 2h**). Spheroid growth remained unperturbed compared with the non-treated negative controls, but we observed gradual development of RFP reporter signals from days 5 to 9, indicating successful EMT induction (**Fig. 2i**). Single-particle MR signals exhibited nearly identical intensity trajectories as those observed in the negative control until day 5, showing progressive compressive stress buildup (**Fig. 2j,k**). However, beyond this point, MR signals reversed toward their original contrast levels, corresponding to a release of the previously accumulated compressive stress by day 9 (**Fig. 2 j,k**). Notably, this reversal coincided with EMT induction, suggesting that stress release occurs through unjamming transitions from a solid-like to a fluid-like state within the spheroids^33^.

## Spatiotemporal stress mapping of developing tumor *in vivo*

Extending the approach beyond engineered spheroids requires testing sensor performance in living tissues with greater mechanical complexity. We therefore evaluated M^3^ sensor performance in brain development of *Xenopus laevis* tadpoles, which undergo rapid morphogenetic and mechanical remodeling and offer optical transparency that enables CFM-MRI investigations throughout organogenesis. Even under these highly dynamic conditions, the M^3^ sensor maintained strong signal fidelity and reproducibility, and MR readouts closely mirrored particle deformation patterns (**Fig. 3a**), remaining consistent with known morphogenetic trajectories of *Xenopus* brain development (**Extended Data Fig. 4**, **Supplementary Note 1**).

**Fig. 3.**
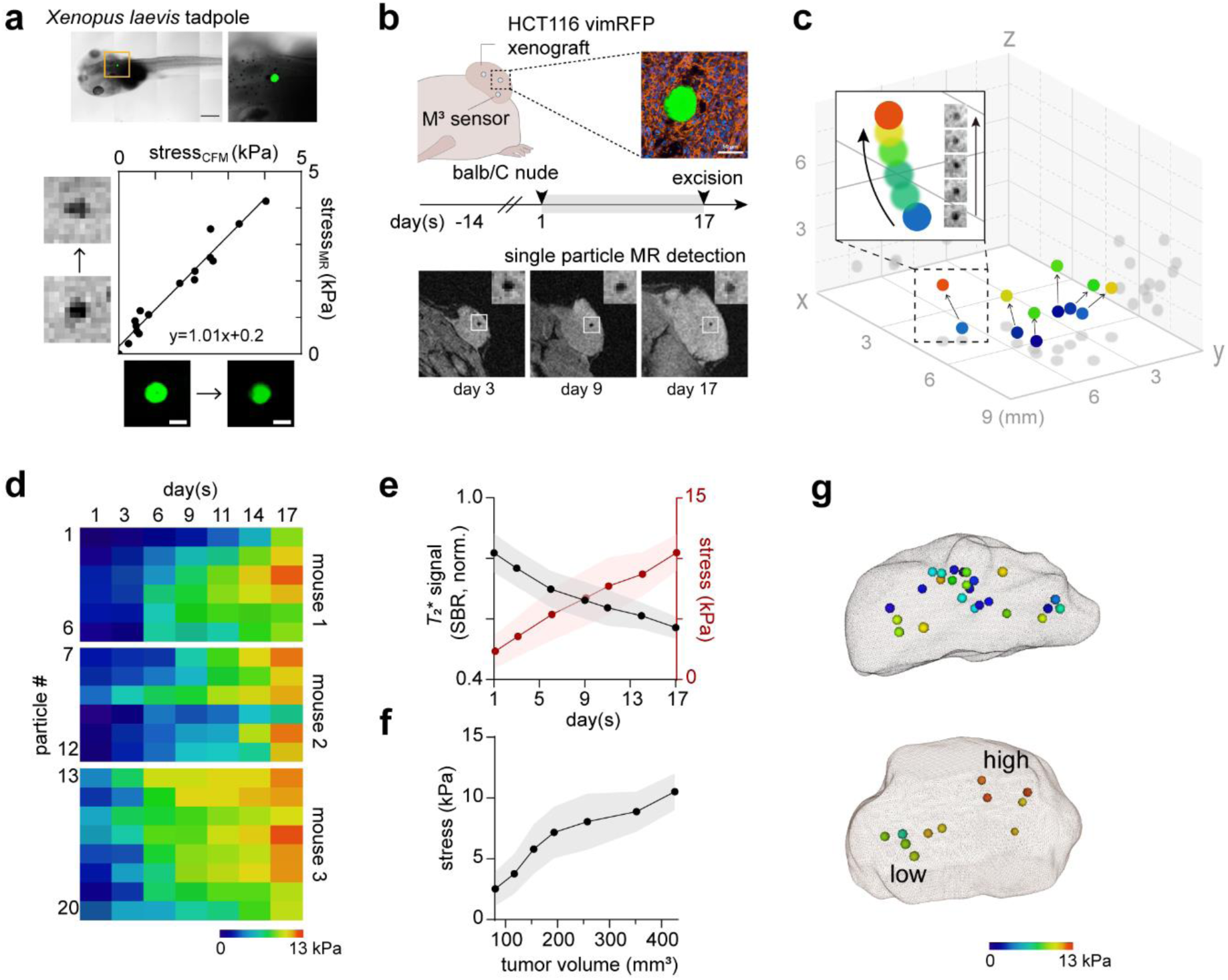
Spatiotemporal stress mapping of developing *in vivo* organisms using M^3^ sensors. **a,** Correlated CFM-MRI investigation of mechanical stress measured with M^3^ sensors. *Xenopus* tadpole showing the implantation site of an M^3^ sensor in hind brain (top) and correlation of CFM-stress measurement with MR-stress measurement, alongside the corresponding CFM and MR image. (bottom, linear correlation with a slope of 1.01; R² = 0.96). **b,** Experimental scheme of stress mapping within developing tumor in a mouse xenograft model. The sensor integration was confirmed by immunohistochemistry (top; green, alginate; blue, DAPI; red, actin). Representative MR images of tumors on day 3, 9, and 17 showing the stress evolution during tumor progression (bottom, inset shows magnified M^3^ sensor). **c,d**, A representative spatiotemporal mapping of dynamic stress development within a tumor (**c**). Representative examples of single-particle stress tracking are shown in panel (**d**). **e,f,** Stress buildup dynamics as a function of time (**e**) and tumor volume (**f**). Data are presented as mean ± s.d. from three mice (*n* = 3; 6-9 sensors per mouse). (**g**) Representative 3D stress dynamics maps in tumors at day 17 highlighting tumor heterogeneity.

Building on these *in vivo* validations, we next applied the M^3^ sensor to a mouse tumor xenograft model to interrogate the spatiotemporal dynamics of mechanical stress in a disease-relevant mammalian context. Tumor development is shaped by diverse factors, including cellular proliferation, matrix remodeling, vascularization, immune infiltration, and cell states, which collectively define the evolving mechanical landscape^34^. To capture these dynamics, we co-injected M^3^ sensors mixed with HCT116 cells subcutaneously, generating xenografts containing sparsely distributed individual particles that were readily resolved by MRI as vivid, dark single-particle signals (**Fig. 3b**). This experimental design enabled longitudinal tracking of local stress changes and 3D spatial mapping of mechanical heterogeneity across different regions of the developing tumor (**Supplementary Fig. 4**). A representative example is shown in **Fig. 3c**, where single-particle M^3^ signals were traced along with their intratumoral positions over a 17-day period. All tracked locations exhibited progressive compressive stress buildup, ranging from 0.5 to 12.4 kPa, but with distinct accumulation kinetics depending on their local microenvironment (**Fig. 3c-f**). We also included stress accumulation maps at day 17 for two additional examples (**Fig. 3g**): one showed a relatively random spatial distribution of stress, whereas another displayed two spatially distinct intratumoral stress regions, highlighting mechanical heterogeneity during tumor growth.

Among the various factors contributing to this mechanical heterogeneity, cell state transitions, particularly EMT, can dramatically remodel the mechanical landscape, as we previously observed in our *in vitro* spheroid models. To investigate whether similar mechanical remodeling accompanies EMT *in vivo*, we generated xenograft tumors from HCT116-vimentin-RFP reporter cells, induced EMT with 5-azacytidine or hepatocyte growth factor (HGF), and monitored local stress dynamics using single-particle M^3^ signals (**Fig. 4a, Supplementary Fig. 5a**). Consistent with our spheroid results, EMT induction led to a reduction in average mechanical stress within the tumor, compared with untreated controls, presumably due to cell state transitions toward a more fluid-like phenotype (**Fig. 4b**). However, unlike the spheroid case that exhibited a narrow deviation of stress values around the mean, we found that stress across the entire EMT-induced tumor was highly heterogeneous, ranging from 3 to 12 kPa (**Fig. 4b**). This increased heterogeneity is likely attributable to spatially uneven EMT induction, as suggested by our analysis of vimentin-RFP reporter distribution, which revealed patchy regions of EMT activation across the tumor (**Fig. 4c**).

**Fig. 4.**
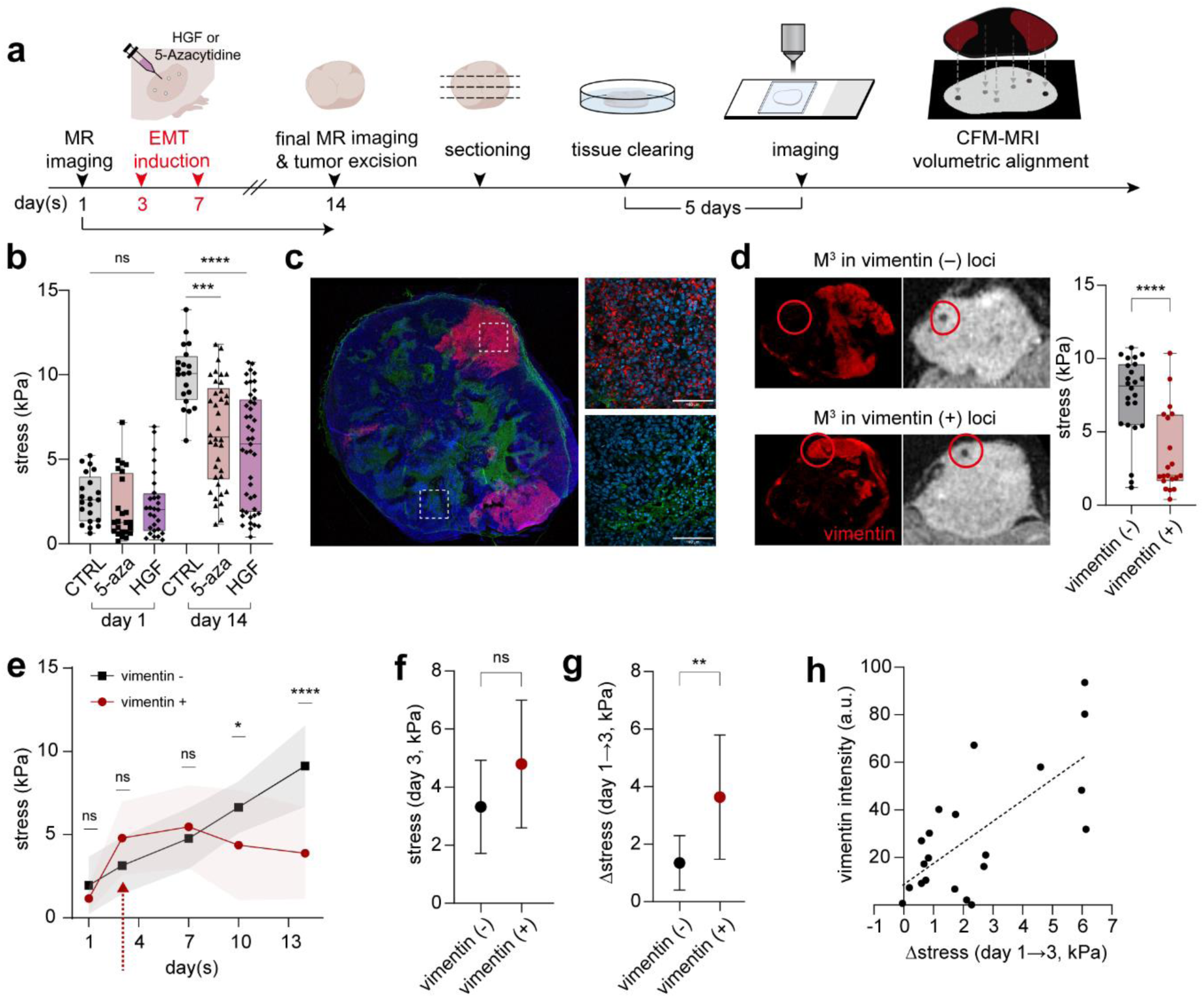
Single particle tracking of local mechanical stress remodeling during EMT. **a**, Schematic showing the workflow of *in vivo* MR imaging and ex vivo tissue CFM imaging, which were subsequently matched for multimodal analysis. **b,** Stress development of different tumor loci before and after EMT-induction. No-treatment control, 5-azacytidine, and HGF groups were measured for comparison. Data were obtained from three independent mice per group (n = 3), with 20-45 sensors analyzed per tumor. **c,** Representative immunofluorescence image of an EMT-induced tumor slice (left) and zoomed-in inset (right) comparing non-EMT and EMT region within the same tissue (blue, DAPI; green, E-cadherin; red, vimentin). **d,** Aligned CFM-MR images showing spatial correspondence between vimentin expression and local stress (left). Stress comparison between vimentin-negative and -positive loci on day 14 (left). **e,** Retrospective stress dynamics of vimentin-negative and -positive loci within tumors over 14 days. **f**,**g,** Comparison of mechanical stress on the day of EMT-induction within loci with the negative and positive signals on the day 14 (**f**) and the corresponding stress change (Δ-stress = day 3→1) (**g**). **h,** Correlation between Δ-stress and vimentin fluorescence intensity showing a positive relationship. Data are presented as mean ± s.d. For box-and-whisker plots, the boxes show the 25th to 75th percentiles, and the whiskers extend to the minima and the maxima. Statistical significance was determined by one-way ANOVA with Tukey’s multiple comparisons test for (**b**), and by two-tailed unpaired Student’s t-test for (**d**), (**e**), (**f**) and (**g**). ****P < 0.0001; ***P<0.001; **P < 0.01; *P < 0.05; ns, non-significant.

To further test this notion, we reconstructed the 3D spatial distribution of vimentin expression across the entire tumor by sequentially imaging 300 µm-thick, SHIELD-cleared^35^ tissue sections, followed by volumetric alignment of the reconstructed fluorescence map with MR tomography data (**Supplementary Fig. 5b, Supplementary Video 2**). We then analyzed the spatial relationship between individual M^3^ sensor locations and vimentin-positive regions within this integrated 3D framework (**Fig. 4d**). M^3^ sensors within the high vimentin expression regions consistently reported lower local stress (3.5 ± 2.8 kPa) compared with those located in vimentin-negative regions (7.4 ± 2.8 kPa), supporting the notion that EMT-rich zones correspond to mechanically more fluid-like microenvironments within the tumor (**Fig. 4d**).

## Critical roles of tissue stress dynamics, rather than static magnitude, in EMT initiation

These findings suggest that EMT-associated mechanical remodeling is reflected in reduced local stress but is also highly spatially heterogeneous, raising the question of what governs selective EMT activation in certain tumor regions while others retain an epithelial phenotype.

To address this, we examined the mechanical microenvironment prior to EMT induction by retrospectively analyzing single-particle M^3^ readouts at each sensor location before treatment (**Fig. 4e**). The average stress level in regions that later underwent EMT was slightly higher than that in EMT-negative regions, but the difference did not reach statistical significance (**Fig. 4f**). In contrast, a far more striking distinction emerged when we compared the stress-loading rate between the two populations (**Fig. 4g,h**). Loci that subsequently transitioned to a mesenchymal state consistently experienced recent, steep increases in local stress, whereas regions with gradual or minimal stress accumulation rarely underwent the transition (**Fig. 4f-h**). These results led us to hypothesize that the dynamics of mechanical stress loading may provide a decisive mechanical cue that primes cells toward EMT.

To test this hypothesis, we returned to the HCT116-vimentin-RFP spheroid system and applied precisely controlled external stresses to directly examine how the temporal dynamics of mechanical loading influence EMT initiation. Spheroids encapsulated within agarose blocks were positioned in a calibrated sandwich compression device and subjected to two distinct dynamic stress regimens (**Fig. 5a,b**). In both cases, the peak compressive stress reached 5 kPa after 48 hours, but the loading profiles differed: one condition increased stress gradually every 6 hours, whereas the other maintained a low baseline stress of 0.5 kPa for 48 hours before rapidly ramping to 5 kPa. These regimens thus delivered either a greater total mechanical input or a greater stress change, respectively, while reaching to identical peak stress. Following this mechanical preconditioning, spheroids were treated with 5-azacytidine, and EMT induction was assessed 24 hours post-treatment – the time point at which EMT was not detectable without external stress application in our previous experiments shown in **Fig. 2i-k** – by quantifying vimentin reporter expression under the two loading regimes. Whereas spheroids exposed to gradual loading or no external stress showed negligible reporter activation, those subjected to acute compressive stress exhibited a markedly increased fluorescence (**Fig. 5a,b**), supporting the notion that the rate of stress application, rather than its absolute magnitude or total load, is the critical determinant of EMT initiation.

**Fig. 5.**
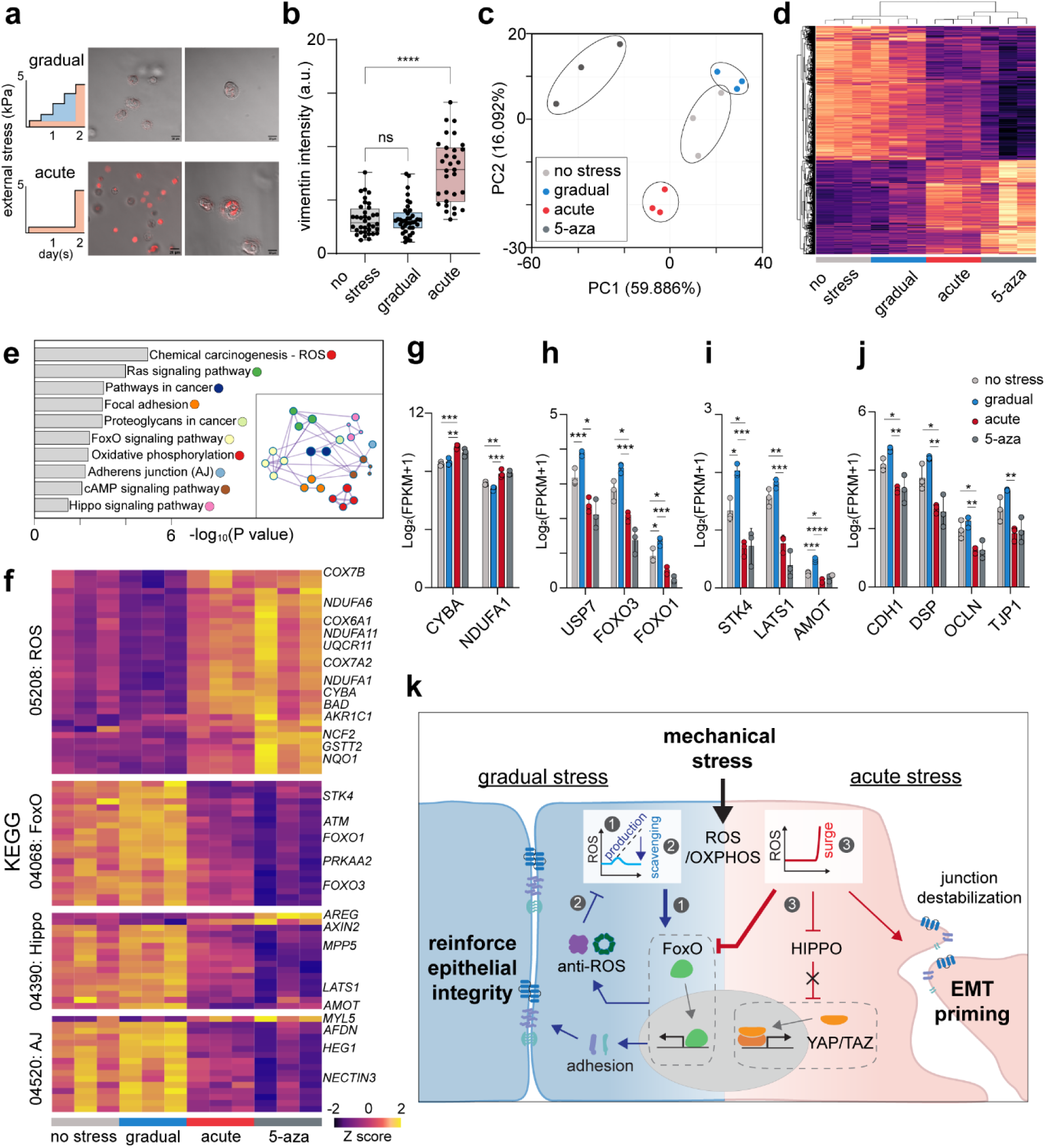
Mechanical stress dynamics direct transcriptional reprogramming during EMT. **a**,**b,** Mechanical loading profiles (gradual and acute stress) applied to tumor spheroids and corresponding CFM images (**a**). Vimentin-RFP quantification (**b**) reveals stronger EMT activation under acute stress than no or gradual stress (*n* = 36 for no stress, *n* = 38 for gradual, and *n* = 33 for acute). **c**,**d,** PCA plot (**c**) and hierarchical clustering heatmap (**d**) of differentially expressed genes (DEGs) of RNA-seq data from spheroids under no stress, gradual stress, acute stress, or 5-azacytidine treated conditions (*n* = 3 per group). Both analyses reveal distinct transcriptomic patterns according to mechanical loading mode. **e,** Significantly enriched KEGG pathways under acute stress (ROS generation, Ras signaling, focal adhesion, proteoglycans in cancer, FoxO signaling, OXPHOS, adherens junction, cAMP, and Hippo signaling). All pathways are interconnected in the gene network map (left) and collectively associated with EMT progression. **f,** Heatmaps of representative pathways - Chemical carcinogenesis-ROS (05208), FoxO (04068), Hippo (04390), adherens junctions (AJ) (04520), highlighting expression changes of genes. **g**-**j,** Comparison of key genes expression values (Log_2_[FPKM+1];FPKM, fragments per kilobase of transcript per million mapped reads) from enriched pathways. (**g**) CYBA and NDUFA1 (ROS/OXPHOS); (**h**) USP7, FOXO3, and FOXO1 (FoxO); (**i**) STK4, LATS1, and AMOT (Hippo); (**j**) CDH1, DSP, OCLN, and TJP1 (AJ). **k,** Proposed model summarizing that how different mechanical-stress dynamics regulate EMT. Under gradual stress, (1) manageable ROS production activates FoxO signaling, (2) promoting antioxidant and adhesion-related gene expression to maintain epithelial integrity and cellular defense. In contrast, Acute stress generates a ROS surge that overwhelms defenses, (3) suppresses FoxO and Hippo signaling, destabilizes junctions, and primes EMT. Data are presented as mean ± s.d. Statistical significance was determined by one-way ANOVA with Tukey’s multiple comparisons test for (**b**) and (**g**-**h**). ****P < 0.0001; ***P<0.001; **P < 0.01; *P < 0.05; ns, non-significant.

## Acute mechanical loading primes EMT-associated transcriptional programs

The observation that acute mechanical loading dramatically enhances EMT induction suggested that such dynamics might initiate transcriptional changes predisposing cells toward mesenchymal reprogramming. To explore this possibility, we performed bulk RNA sequencing of spheroids subjected to either gradual or acute compressive loading and compared their transcriptional landscapes with those of untreated controls (no stress, epithelial phenotype) and 5-azacytidine-treated spheroids (positive control, mesenchymal phenotype) (n=3 per condition). Principal component analysis (PCA) revealed clear separation among the four experimental groups, indicating high reproducibility and distinct transcriptional states (**Fig. 5c**). Spheroids exposed to gradual stress loading clustered closely with the unstressed controls, exhibiting only modest transcriptional deviations relative to epithelial baseline (**Fig. 5c**). In contrast, those subjected to acute stress loading positioned intermediately along PC1 between the negative and positive controls, suggesting a transitional transcriptional state under this condition. Hierarchical clustering further supported this pattern: spheroids under gradual loading retained epithelial-like transcriptomic signatures, whereas those under acute loading shifted toward mesenchymal-like profiles (**Fig. 5d, Extended Data Fig. 5a-d**).

Pathway enrichment analysis revealed that acute stress loading activated a distinct set of molecular pathways, including reactive oxygen species (ROS) generation, Ras signaling, focal adhesion, proteoglycans in cancer, Forkhead box O (FoxO) signaling, oxidative phosphorylation (OXPHOS), adherens junctions, cAMP signaling, and Hippo signaling – all of which are closely linked to EMT induction and progression^36–40^ (**Fig. 5e, Extended Data Fig. 5e**). Specifically, acute mechanical stress triggered a coordinated transcriptional reprogramming marked by activation of a NOX2 (e.g., CYBA)-mitochondrial ROS/OXPHOS (e.g., COX, NDUF, UQCR, ATP5) axis, coupled with suppression of antioxidative pathways (e.g., FOXO, AMPK, ATM) and broad attenuation of growth factor/RAS-PI3K signaling (e.g., IGF1R, SOS1, KRAS) (**Fig. 5f-h, Extended Data Fig. 5f**). Concomitantly, Hippo pathway brakes (e.g., STK4, LATS1, AMOT) were reduced (**Fig. 5f,i**). Transcripts associated with extracellular matrix production, glycocalyx biosynthesis, and cell-cell junctions were broadly downregulated, whereas actomyosin contractility was sustained via reduced myosin phosphatase expression (PPP1R12B) (**Fig. 5f,j** and **Extended Data Fig. 5f**). All these changes converge to stabilize a state favoring EMT and enhanced migratory capacity. In contrast, spheroids subjected to gradual mechanical stress displayed modest but inverse transcriptional trends across these pathways (**Fig. 5f-j**). Particularly, NOX2-mitocondrial ROS/OXPHOS genes were downregulated, whereas antioxidative and FOXO-dependent programs were upregulated (**Fig. 5f-h**). These results suggest that cells exposed to sustained, manageable stress activate protective antioxidative responses that reinforce epithelial stability, whereas acute stress overwhelms these defense mechanisms shifting the balance toward ROS amplification, energy remodeling, and mesenchymal transition (**Fig. 5k**).

## Discussion

A central question in mechanobiology is how the magnitude, timing, and spatial pattern of mechanical stress collectively regulate cellular and tissue behavior. Although decades of work have established that mechanical cues guide development, regeneration, and disease progression, progress toward a quantitative understanding has been limited by the absence of methods that directly report stress within living tissues. The M³ sensor presented here overcomes this limitation through a simple yet powerful mechanoMR transduction mechanism, enabling spatiotemporal mapping of tissue stress across scales previously inaccessible to mechanobiology. This capability transforms how dynamic tissue mechanics can be visualized and interrogated *in vivo*. The EMT model illustrates this power, revealing how the magnitude, rate, and spatial distribution of stress collectively dictate cellular perception, transcriptional reprogramming, and phenotypic transition. Our finding – that the time derivative of local mechanical stress, rather than its magnitude, determines whether cells engage cytoprotective or reprogramming pathways – was possible only through direct, *in vivo* quantification of stress dynamics.

Beyond this specific cancer model, this technology opens a quantitative window into developmental and pathological processes governed by evolving mechanical landscapes. Stress-mapping with M³ sensors could reveal how forces sculpt morphogenesis, tissue repair, fibrosis, cardiovascular remodeling, and neurodegeneration – contexts where the timing and amplitude of stress are thought to instruct cellular fate but remain experimentally elusive. By coupling MR anatomic imaging with embedded soft-matter probes, M³ sensors bridge the gap between molecular imaging and tissue mechanics, providing a physical map of the states that shape biological form and function.

Mechanistically, our results also show that acute stress loading primes, but does not fully induce EMT. What does differentiate this primed state from complete mesenchymal conversion? Transcriptomic comparisons between acutely stressed and chemically induced EMT revealed that mechanical stress activates EMT-associated transcriptional programs but fails to engage the chemokine and TNF feedback loops required for stable mesenchymal commitment^41,42^ (**Extended Data Fig. 6**). Notably, HMGB2, a recently identified nuclear compression-induced EMT driver in a skin cancer model – remained unchanged under both mechanical and chemical stimulation (**Extended Data Fig. 7**), suggesting that this pathway depends on the extreme nuclear deformation by very stiff tumor environments like skin cancer^43^. We propose that, in softer epithelial tissues, transient stress elevations act as mechanical primers, sensitizing cells to biochemical EMT cues without large-scale nuclear deformation, defining an alternative, HMGB2-independent route to phenotypic plasticity.

In summary, the M³ platform provides a quantitative framework for decoding how tissues generate and interpret mechanical forces *in vivo*. By allowing direct visualization of stress evolution from the cellular to tissue scale, it redefines how mechanical information can be captured and connected to biological outcomes. Continued technical refinement, such as miniaturization to nanoscale for systemic delivery and logical probe design that distinguishes compressive from shear stresses, will further extend its reach from basic mechanobiology to translational applications, where mechanical states may emerge as diagnostic or prognostic indicators of disease.

## Methods

### Synthesis of functionalized MA-alginate

Methacrylated alginate (MA-alginate) was prepared following a previously reported protocol with minor adjustments^44^.

#### DBCO-MA-alginate.

40 mg of MA-alginate was dissolved in 9 mL of distilled water. Prepare 160 μL of 100 mM DBCO-PEG_4_-amine (Broadpharm) dissolved in dimethyl sulfoxide (DMSO) and 19.2 μmol of 4-(4,6-dimethoxy-1,3,5-triazin-2-yl)-4-methyl-morpholinium chloride (DMTMM, Sigma-Aldrich) in DMSO. The two solutions were added to the MA-alginate solution to obtain a final reaction volume of 10 mL and incubated under continuous shaking overnight. The product was dialyzed using 15 kDa dialysis tubing (Fisher Scientific) for 3 days and subsequently freeze-dried under vacuum.

#### Fluorescein-MA-alginate.

8.34 mg of fluoresceinamine (isomer Ⅰ, Sigma-Aldrich) dissolved in 500 μL DMSO was used in place of DBCO-PEG_4_-amine, and the reaction was performed using the same procedure.

#### cRGD-MA-alginate.

0.1g of MA-alginate, 25.8 μmol of 1-ethyl-3-[3-dimethylaminopropyl]carbodiimide hydrochloride (EDC, Thermo Scientific), and 51.6 μmol sulfo-N-hydroxysulfosuccinimide (sulfo-NHS, Sigma-Aldrich) were dissolved in phosphate buffer (pH 7.2) and mixed with cyclo(-Arg-Gly-Asp-D-Phe-Lys) (cyclo(-RGDfK , ≥95%, Anaspec) to a final concentration 150 μM. The reaction mixture (10 mL total) was incubated for 20 h, dialyzed for 3 days, and freeze-dried under vacuum.

### Synthesis of M^3^ sensors

For preparing azide-functionalized MNPs, Zinc-doped iron oxide (Zn_0.4_Fe_2.6_O_4_) magnetic nanoparticles (MNPs) were synthesized^25^ and subsequently silica-coated and PEGylated according to established protocols described previously^24^.

Microfluidic devices were produced using conventional photolithography and PDMS replica molding, following well-established procedures. Detailed fabrication procedures are provided in the Supplementary Methods.

To synthesize M^3^ sensors, a pre-gel mixture was prepared by combining 40 µl of DBCO-MA-alginate (2% w/v in distilled water), 40 µl of cRGD-MA-alginate (2% w/v in distilled water), 20 µl of Fluorescein-MA-alginate (2% w/v in distilled water), and 10 µl of poly(ethylene glycol) diacrylate (PEGDA 700, Sigma-Aldrich). MNPs were added to the mixture at a final concentration of 1500 nM. For photopolymerization, 40 µl of lithium phenyl-2,4,6-trimethylbenzoylphosphinate (LAP, 2.5 % w/v in distilled water, Sigma-Aldrich) was added, and distilled water was used to adjust the final volume to 200 µl. The pre-gel solution was injected into the microfluidic device and emulsified into droplets using a continuous phase of fluorinated oil (HFE-7500, Kemis) supplemented with 2% (w/v) 008-Fluorosurfactant (RAN Biotechnologies). Flow rates were set to 800 µl h^−1^ for the continuous phase and 20 µl h^−1^ for the dispersed phases using syringe pump (Harvard Apparatus). For UV-initiated polymerization, emulsions were irradiated with a 365 nm UV-light source (Hamamatsu Photonics, 800 mW/cm^-2^) within the channel and collected into Eppendorf tube. Droplets containing polymerized M^3^ sensors were broken using 20% (v/v) 1H,1H,2H,2H-perfluoro−1-octanol (PFO, Sigma-Aldrich). The recovered M^3^ sensors were subsequently washed several times with PBS to remove residual non-crosslinked monomers and oil contaminants.

### Single particle MR-stress calibration

#### Sandwich device fabrication.

The sandwich device was fabricated exclusively from non-metallic materials to ensure compatibility with MRI. Using a laser cutter, cut two acrylic plates (21 × 60 × 2 mm) and make four holes (4 mm diameter). Prepare four plastic screws and eight nuts to connect the two acrylic plates, allowing one to be fixed below and the other to move up and down. Cylindrical agarose template (diameter & height 10 mm) was customed by 3D printer (Ultimaker Cura 3.0). For MR imaging, M^3^ sensors were mixed with agarose (500 particles/mL) and injected into cylindrical template just before solidification. Agarose was solidified in cylindrical template at 25 °C, at the desired concentration (0.5, 1, 2 %). Place an agarose block in the middle of the sandwich device and adjust the calculated z-displacement to apply the desired stress by compression. Press the agarose block and fasten using four nuts while applying stress, and after 10 minutes, perform MRI imaging.

#### Finite element modeling.

To quantify the stress caused by displacement on a M^3^ sensors-embedding agarose gel system, three-dimensional finite element models were constructed using ABAQUS (Standard). The non-linear mechanical behavior of the system was modeled as a Neo-Hookean material, with Lamé parameters derived from the shear modulus and Poisson’s ratio. The values used were Poisson’s ratio ν=0.31 and shear modulus G=0.4, 1.5, and 4.7 kPa for agarose concentrations (w/v) of 0.5%, 1.0%, and 2.0%, respectively. The geometric representation of the system was simplified to a cylinder, 10 mm in diameter and height and subjected to compression via displacement applied to its top surface. The stress distribution within the system was uniform and was directly extracted from the simulation results. The model was discretized using solid linear brick elements (C3D8, verification needed).

#### In vitro MR Imaging.

For imaging, the phantom was submerged in PBS and MRI was performed on a 3 T preclinical MRI system (MRS 3017 model, MR Solutions) with the following specifications: clear-bore size=17 cm; gradient strength=600mTm^−1^ ; radiofrequency amplifier power=500W; radiofrequency coil=mouse body volume coil; and operating software=Preclinical Scan. To optimize the visualization of stress-dependent signal alterations, MRI data were acquired using a Fast Low Angle Shot (FLASH) magnetic resonance imaging sequence, which provide high sensitivity to *T*_2_ contrast while maintaining rapid acquisition. Other imaging parameters were: repetition time 35 ms; echo time = 20 ms; flip angle=12.0 °; field of view (FOV) = 25.6 × 51.2 mm; voxel size = 0.1 × 0.1 × 0.2 mM^3^. Scan times were 35 min. Magnetic resonance images were analyzed using open-source software (ImageJ). Regions of interest (ROIs) were defined to capture the *T*_2_*-weighted gray intensity from the phantom or spheroid region on MR images. All experiments had the same ROI dimensions (3 × 3 pixels). The mean gray intensity was calculated for each ROI. The M^3^ sensor signal was obtained by setting a ROI at the center of strong intensity in both upper and lower images and calculating the average value. The background signal was determined by measuring the signal intensity of agarose or Matrigel at four non-particle areas diagonally adjacent to the M^3^ sensor and calculating the average value. To obtain a value proportional to the relaxation rate, the reciprocal of each signal was used in the calculation.

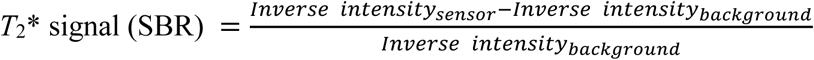

#### Calibration of MR signal to mechanical stress.

To calibrate the relationship between stress (S) and the *T*_2_* signal (SBR), agarose block containing M^3^ sensors was imaged under a series of known stress levels. The *T*_2_* signal value at each applied stress was plotted and fitted by nonlinear least-squares to an empirical bi-exponential decay model, *T*_2_* signal = 0.1344 e^−0.926S^ + 0.864 e^−0.0405S^. This calibration curve was subsequently used estimate local stress from measured *T*_2_* signal. For each *T*_2_* signal, the corresponding stress S was obtained by numerically solving the above equation for S using the Newton-Raphson method.

### Cell culture

HCT116 vimRFP cell line was purchased from ATCC (CCL-247EMT) and grown in McCoy’s medium (Gibco). A549 cell lines and MRC-5 cell lines were cultured in DMEM medium (Gibco), and MDA-MB-231 cell lines were cultured in RPMI-1640 medium (Gibco). All mediums were supplemented with 10 % heat-inactivated FBS (Life Technologies) and 1 % Pen-Strep (Life Technologies). All cell lines were maintained at 37 ℃ in a humidified incubator with 5 % CO2 and passaged every 2-3 days, depending on confluency using 0.25 % trypsin-EDTA (Gibco). To prevent mycoplasma contamination, cells were routinely maintained in growth medium supplemented with a mycoplasma removal agent (500:1 dilution; MPbio). The reagent was included at every medium change throughout culture.

### Quantification of mechanical stress in tumor spheroids

Spheroids embedded M^3^ sensor were generated using sacrificial gelatin microwells, fabricated using a PDMS stamp, as previously described^45^ with minor modifications. Briefly, dissociated cells were mixed with M^3^ sensors and centrifuged into gelatin microwells to induce aggregation, followed by transfer into Matrigel for long-term culture. Detailed procedures are provided in the Supplementary Methods. In the fibrotic spheroid experiment, Nintedanib (Sigma-Aldrich) was treated at a concentration of 1 μM in DMEM medium and medium changes performed every 2 days to inhibit fibrosis. For EMT spheroid experiment, 5-azacytidine (Sigma-Aldrich) was prepared at 10 mM concentration in DMSO. 5-azacytidine was treated at a concentration of 10 μM in McCoy’s medium, with medium changes conducted every 2 days to induce EMT. MR imaging and stress quantification of spheroids were performed using the same calibration curve and in vitro MRI acquisition protocols described above.

### *In vivo* stress measurement

#### Mouse xenografts.

5-week old male immune-deficient Balb/C nude mice (CAnN.Cg-Foxn1 nu/CrlOri, Orient Bio.) All mice were maintained under the approved by the Committee on the Ethics of Animal Experiments of Yonsei University (IACUC-202307-1705-02). HCT116 vimRFP (CCL-247EMT, ATCC) cells were used to establish tumors for *in vivo* experiments. The viable cells were counted after Trypan blue staining. 5 × 10^6^ cells were counted, washed in PBS three times, resuspended in 100 µl of PBS containing approximately 100 M^3^ particles. The mixture was then subcutaneously grafted into mouse hind limb. MRI monitoring was started when tumor was grown until it reached a size over 60 mM^3^.

#### MR acquisition.

For MRI scanning, mice were anaesthetized with isoflurane (Hana Pharm) in oxygen. The dosage of isoflurane was 3% for induction and 1.5 ∼ 2% for maintenance. The dosage was controlled using an anesthesia system (L.M.S Korea). The MR imaging started when tumor was grown a size over 60 mM^3^ and imaging was conducted every 2-3 days to monitor the tumor progression during 17 days. Images were obtained using Fast Low Angle Shot (FLASH) magnetic resonance imaging sequence. Other imaging parameters were: repetition time 35 ms; echo time = 20 ms; flip angle=12.0 °; field of view (FOV) = 25.6 × 51.2 mm; voxel size = 0.1 × 0.1 × 0.2 mM^3^. Scan times were 70 min. *T*_2_* signals were calculated in the same manner as in the *in vitro* MR imaging experiments. Although M^3^ sensors were mixed with cells to form tumors while minimizing aggregation, particle aggregation could still occur under *in vivo* conditions. Based on the *in vitro* calibration results, aggregated particles were identified and excluded from the analysis if their *T*_2_* signal exceeded 2.5 and if the signal covered a contiguous area larger than a 3 × 3 pixels region. Only single, non-aggregated particles were included for quantitative analysis.

#### EMT induction.

For the EMT study, Hepatocyte Growth Factor human (HGF, Sigma-Aldrich, H9661) diluted to 10 ug/mL in PBS containing 0.1% bovine serum albumin (BSA). When tumor size exceeded 100 mM^3^, intratumoral injection of 30 µg/kg HGF was performed, with a second injection administered 3 days later. The tumor was monitored by MRI during 14 days.

### Volumetric alignment of the reconstructed CFM images with MRI

#### Ex vivo CFM imaging for 3D reconstruction.

Embedded tumors were sectioned by 300 μm thickness using a microtome (Leica Microsystems GmbH). The tumor slices were incubated in SHIELD-OFF solution (ice-cold PBS containing supernatant of 10 % GE38) at cold room for overnight. After SHIELD-OFF step, the tumor slices were placed in the SHIELD-ON solution (0.1 M sodium carbonate buffer at pH 10 containing supernatant of 1 % GE38, prewarmed to 37 °C) and incubated at 37 °C for 3 h. SHIELD-fixed tumor slices were cleared through delipidation step for 2 days in SDS-based clearing buffer (300 mM SDS, 10 mM sodium borate, 100 mM sodium sulfite, pH 9.0). Delipidated tumor slices were incubated in optical clearing solution (EasyIndex, lifecanvastechnologies) until the tissue became transparent and imaged vimentin-RFP fluorescence using an upright confocal microscope (Leica Microsystems GmbH).

#### Volumetric alignment.

MRI scans and CFM images were segmented in 3D Slicer, and the segmented volumes were exported as STL surface meshes. M^3^ sensors were manually identified within the MRI volume and labeled in 3D Slicer to extract their 3D coordinates. Both the STL surface meshes and M^3^ sensor coordinates were then imported into MATLAB for alignment and downstream analysis. Using custom MATLAB scripts (available at https://github.com/JunLab-UCSF/M3_VolumeRegistration), the MRI-derived and CFM-derived meshes were first translated to a common origin and then aligned using non-rigid coherent point drift (pcregistercpd). The resulting transformation was applied to the M^3^ sensor coordinates to map their positions into the CFM reference frame. These registered particle locations were subsequently used for spatial analyses.

### Immunohistochemistry (IHC)

For immunohistochemical analysis, anesthetized animals were perfused with PBS and 4% paraformaldehyde (Thermo Fisher Scientific) followed by excising Tumors. Tumors were stored at -80°C after embedded in peel-a-way embedding mold (Sigma-Aldrich) using frozen sectioning compound (FSC 22 Clear, Surgipath). The frozen tissue blocks were sectioned into 8 μm thickness using a cryocut microtome (Leica Microsystems GmbH, Germany) previously fluorescence staining. The sections after washing with PBS, the tissues were fixed with 4% paraformaldehyde, followed by treatment with 0.1% TritonX-100 and SuperBlock Blocking Buffer (SuperBlock (PBS), Thermo Fisher Scientific) to permeabilization and prevent nonspecific binding. Subsequently primary antibodies (anti-E-cadherin (Invitrogen, MA5-12547, 1:200) was applied and incubated overnight at 4°C. The following day, the tissues were washed with a 0.1% Tween-20 PBS solution. Then, secondary antibodies (anti-mouse IgG (Alexa Fluor 488, Abcam, 1:300) were applied for 1 hour at room temperature. In the case of Rhodamine Phalloidin (R415, Invitrogen 1:300); was diluted with a 0.1% Tween-20 SuperBlock solution. Additionally, NucBlue™ Live ReadyProbes™ Reagent (Hoechst 33342, Thermo Fisher Scientific) was stained under the same conditions for 30 minutes. After PBS washing, the stained tissues were mounted with Crystal Mount™ Aqueous Mounting Medium (Sigma-Aldrich). Fluorescence signals were observed using a confocal microscope (Zeiss, Germany) equipped with ZEN software and analyzed using ImageJ.

### Transcriptomic analysis of tumor spheroids

#### Tumor spheroids compression.

To form cells embedded agarose matrix, dissolve low melting point agarose (UltraPure™ LMP Agarose, Invitrogen) in PBS at 2%, then cool it down to 37 °C. HCT116 tumor cells were dissociated from culture plates, resuspended in 2% LMP agarose at a concentration of 10^7^ cells/mL. The mixture was injected into a cylindrical template and solidified at 25 °C. Allow the cells to form a spheroid within the agarose for 3 days, then compress the spheroid-agarose matrix using the sandwich device from the previous experiment. To apply gradual stress, the spheroid-embedded agarose was compressed by 0.2 mm, followed by five additional compression steps of 0.4 mm each over 2 days. For acute stress, another agarose sample was initially compressed by 0.2 mm, followed by an acute displacement of 2 mm at the time point corresponding to the sixth compression of the gradual group, ensuring both groups reached the same final displacement. After compression, both samples were incubated overnight. The levels of vimentin-RFP expression were subsequently assess using a Zeiss confocal microscope for comparative analysis.

#### RNA sequencing preparation.

After confocal imaging, RNA sequencing analysis was performed to compare transcriptomic profiles of tumor spheroids cultured under acute and gradual stress conditions. To isolate cells from the agarose matrix following overnight compression, the agarose gel was finely chopped and transferred into 970 μL of PBS. Then, 30 μL of agarase (Thermo Scientific) was added, and the mixture was incubated at 37 °C for 1 hour to enzymatically degrade the agarose. After incubation, the samples were centrifuged, and the cell pellet was washed 3 times with PBS to remove residual enzyme. The final pellet was resuspended in TRIzol reagent (QIAzol Lysis Reagent, QIAGEN) and stored at −80 °C for subsequent RNA extraction. Total RNA extraction, library preparation and sequencing were performed by Macrogen (Seoul, Korea), as detailed in the Supplementary Methods.

#### Differential gene expression analysis.

Differential gene expression (DEG) analysis was performed by Macrogen (Seoul, Korea) using DESeq2 v 1.38.3 with raw counts as input. Principal component analysis (PCA) plot was generated to assess expression similarity among samples. Statistical significance of differential expression was determined using DESeq2 nbinom WaldTest and fold change and p-value were extracted from the WaldTest results. All p-values were adjusted using the Benjamini-Hochberg algorithm to control false discovery rate (FDR). Hierarchical clustering on rlog transformed values for significant genes was performed using Euclidean distance and complete linkage. Gene-enrichment and functional annotation analysis were carried out using in-house KEGG Viewer script against KEGG pathway database. Adjusted p-values were derived from a two-sided modified Fisher’s exact test with Benjamini-Hochberg correction. All data analysis and visualization of differentially expressed genes were conducted using R 4.2.2 and the Metascape web-based platform.

### Statistical analyses

Statistical analysis was performed in GraphPad Prism 10.0 (GraphPad). Figure legends indicate all statistical test used in the figure. Unless otherwise noted in the figure legends, statistical differences were determined using Student’s t-test (two-tailed unpaired or paired t-test, depending on the experiment) when only two groups were compared or by ordinary one-way ANOVA flowed by Tukey post-hoc test when multiple groups were analyzed. The number of samples (‘n’) used for each experimental analysis is reported in the figure legends. No statistical methods were used to predetermine sample size, and experiments were not randomized. All samples used in each set of experiments were equal, except the experimental condition being tested. All experiments were performed with appropriate control. The investigators were not blinded to sample allocation during experiments or outcome assessment.

### Data availability

The statistical data are provided with the paper as source data. All other data supporting the findings are available within the article and its Supplementary Information, or from the corresponding authors upon reasonable request.

## Acknowledgement

This work was supported by the National Institute of Health and the National Institute of General Medical Science (NIGMS, R35GM134948) (Y.J.), The Sandler Program for Breakthrough Biomedical (PBBR) Research which is partially funded by the Sandler Foundation (Y.J.), and the Institute for Basic Science (IBS-R026-D1) (J.C.). We thank Drs. Abate (UCSF) for consultation on microfluidic synthesis, Lungerich (IBS) for rheometer access, Kim (Yonsei U) and Sungwhi Kang (IBS) for consultation on MRI analysis.

## Author contribution

M.A., Y.J. and J.C. conceived and designed the project. M.A. performed overall experiments and coordinated the study. R.Y. conducted the *in vivo Xenopus* tadpole and mouse imaging experiments. A.L. analyzed and interpreted data related to 3D image reconstruction. J.J. fabricated microfluidic device and synthesized PEGDA microparticles. J.O. and H.J. performed *Xenopus* tadpole microinjection experiments. Y.K. carried out the agarose compression-stress simulation. M.K. contributed to the project discussion. Y.J. and J.C. supervised all aspects of the project. M.A., Y.J. and J.C. wrote the manuscript.

## Competing interests

The authors declare no competing interests.

## Additional information

**Correspondence and requests for materials** should be addressed to Young-wook Jun or Jinwoo Cheon.

## Notes

### Competing Interest Statement

The authors have declared no competing interest.

## References

1 Brugués, A. et al. Forces driving epithelial wound healing. Nat. Phys. 10, 683–690 (2014).

2 Northey, J. J., Przybyla, L. & Weaver, V. M. Tissue force programs cell fate and tumor aggression. Cancer Discov. 7, 1224–1237 (2017).

3 Mammoto, T. & Ingber, D. E. Mechanical control of tissue and organ development. Development 137, 1407–1420 (2010).

4 Hahn, C. & Schwartz, M. A. Mechanotransduction in vascular physiology and atherogenesis. Nat. Rev. Mol. Cell Biol. 10, 53–62 (2009).

5 Paszek, M. J. et al. Tensional homeostasis and the malignant phenotype. Cancer cell 8, 241–254 (2005).

6 Binnig, G., Quate, C. F. & Gerber, C. Atomic force microscope. Phys. Rev. Lett. 56, 930 (1986).

7 Neuman, K. C. & Nagy, A. Single-molecule force spectroscopy: optical tweezers, magnetic tweezers and atomic force microscopy. Nat. Methods 5, 491–505 (2008).

8 Hutson, M. S. et al. Forces for morphogenesis investigated with laser microsurgery and quantitative modeling. Science 300, 145–149 (2003).

9 Etournay, R. et al. Interplay of cell dynamics and epithelial tension during morphogenesis of the Drosophila pupal wing. Elife 4, e07090 (2015).

10 Vorselen, D. et al. Microparticle traction force microscopy reveals subcellular force exertion patterns in immune cell–target interactions. Nat. Commun. 11, 20 (2020).

11 Grashoff, C. et al. Measuring mechanical tension across vinculin reveals regulation of focal adhesion dynamics. Nature 466, 263–266 (2010).

12 Stabley, D. R., Jurchenko, C., Marshall, S. S. & Salaita, K. S. Visualizing mechanical tension across membrane receptors with a fluorescent sensor. Nat. Methods 9, 64–67 (2012).

13 Wang, X. & Ha, T. Defining single molecular forces required to activate integrin and notch signaling. Science 340, 991–994 (2013).

14 Nia, H. T. et al. Solid stress and elastic energy as measures of tumour mechanopathology. *Nat*. Biomed. Eng. 1, 0004 (2016).

15 Larin, K. V. & Sampson, D. D. Optical coherence elastography–OCT at work in tissue biomechanics. Biomed. Opt. Express 8, 1172–1202 (2017).

16 Mueller, S. & Sandrin, L. Liver stiffness: a novel parameter for the diagnosis of liver disease. Hepat. Med., 49–67 (2010).

17 Campàs, O. et al. Quantifying cell-generated mechanical forces within living embryonic tissues. Nat. Methods 11, 183–189 (2014).

18 Mohagheghian, E. et al. Quantifying compressive forces between living cell layers and within tissues using elastic round microgels. Nat. Commun. 9, 1878 (2018).

19 Dolega, M. E. et al. Cell-like pressure sensors reveal increase of mechanical stress towards the core of multicellular spheroids under compression. Nat. Commun. 8, 14056 (2017).

20 Lee, W. et al. Dispersible hydrogel force sensors reveal patterns of solid mechanical stress in multicellular spheroid cultures. Nat. Commun. 10, 144 (2019).

21 Zhang, S. et al. Intravital measurements of solid stresses in tumours reveal length-scale and microenvironmentally dependent force transmission. *Nat*. Biomed. Eng. 7, 1473–1492 (2023).

22 Seano, G. et al. Solid stress in brain tumours causes neuronal loss and neurological dysfunction and can be reversed by lithium. *Nat*. Biomed. Eng. 3, 230–245 (2019).

23 Serwane, F. et al. In vivo quantification of spatially varying mechanical properties in developing tissues. Nat. Methods 14, 181–186 (2017).

24 Kim, W.-S. et al. Magneto-acoustic protein nanostructures for non-invasive imaging of tissue mechanics in vivo. Nat. Mater. 23, 290–300 (2024).

25 Jang, J. t., et al. Critical enhancements of MRI contrast and hyperthermic effects by dopant-controlled magnetic nanoparticles. Angew. Chem. Int. Ed. 48, 1234–1238 (2009).

26 Mongera, A. et al. A fluid-to-solid jamming transition underlies vertebrate body axis elongation. Nature 561, 401–405 (2018).

27 Maniou, E. et al. Quantifying mechanical forces during vertebrate morphogenesis. Nat. Mater. 23, 1575–1581 (2024).

28 Paquet, C. et al. Clusters of superparamagnetic iron oxide nanoparticles encapsulated in a hydrogel: a particle architecture generating a synergistic enhancement of the T2 relaxation. ACS Nano 5, 3104–3112 (2011).

29 Tevis, K. M., Colson, Y. L. & Grinstaff, M. W. Embedded spheroids as models of the cancer microenvironment. Adv. Biosyst. 1, 1700083 (2017).

30 Montel, F. et al. Stress clamp experiments on multicellular tumor spheroids. Biophys. J. 102, 220a (2012).

31 Varol, R. et al. Acousto-holographic reconstruction of whole-cell stiffness maps. Nat. Commun. 13, 7351 (2022).

32 Cao, D. et al. 5-Azacytidine promotes invadopodia formation and tumor metastasis through the upregulation of PI3K in ovarian cancer cells. Oncotarget 8, 60173 (2017).

33 Coban, B., Bergonzini, C., Zweemer, A. J. & Danen, E. H. Metastasis: crosstalk between tissue mechanics and tumour cell plasticity. Br. J. Cancer 124, 49–57 (2021).

34 Nia, H. T., Munn, L. L. & Jain, R. K. Physical traits of cancer. Science 370, eaaz0868 (2020).

35 Park, Y.-G. et al. Protection of tissue physicochemical properties using polyfunctional crosslinkers. Nat. Biotechnol. 37, 73–83 (2019).

36 Cheung, E. C. & Vousden, K. H. The role of ROS in tumour development and progression. Nat. Rev. Cancer 22, 280–297 (2022).

37 Tripathi, K. & Garg, M. Mechanistic regulation of epithelial-to-mesenchymal transition through RAS signaling pathway and therapeutic implications in human cancer. J. Cell Commun. Signal. 12, 513–527 (2018).

38 Jiramongkol, Y. & Lam, E. W.-F. FOXO transcription factor family in cancer and metastasis. Cancer Metastasis Rev. 39, 681–709 (2020).

39 Le Bras, G. F., Taubenslag, K. J. & Andl, C. D. The regulation of cell-cell adhesion during epithelial-mesenchymal transition, motility and tumor progression. Cell Adhes. Migr. 6, 365–373 (2012).

40 Dey, A., Varelas, X. & Guan, K.-L. Targeting the Hippo pathway in cancer, fibrosis, wound healing and regenerative medicine. Nat. Rev. Drug Discov. 19, 480–494 (2020).

41 Su, S. et al. A positive feedback loop between mesenchymal-like cancer cells and macrophages is essential to breast cancer metastasis. Cancer cell 25, 605–620 (2014).

42 Yi, M. et al. Targeting cytokine and chemokine signaling pathways for cancer therapy. Signal Transduct. Target. Ther. 9, 176 (2024).

## References

43 Hunter, M. V. et al. Mechanical confinement governs phenotypic plasticity in melanoma. Nature, 1–11 (2025).

44 Wang, X., Hao, T., Qu, J., Wang, C. & Chen, H. Synthesis of thermal polymerizable alginate-GMA hydrogel for cell encapsulation. J. Nanomater. 2015, 970619 (2015).

45 Cerchiari, A. et al. Formation of spatially and geometrically controlled three-dimensional tissues in soft gels by sacrificial micromolding. Tissue Eng. Part C Methods 21, 541–547 (2015).

