## Extended data figures for "MechanoMR microparticle (M^3^) sensors reveal dynamic stress loading as a driver of epithelial-mesenchymal transition"

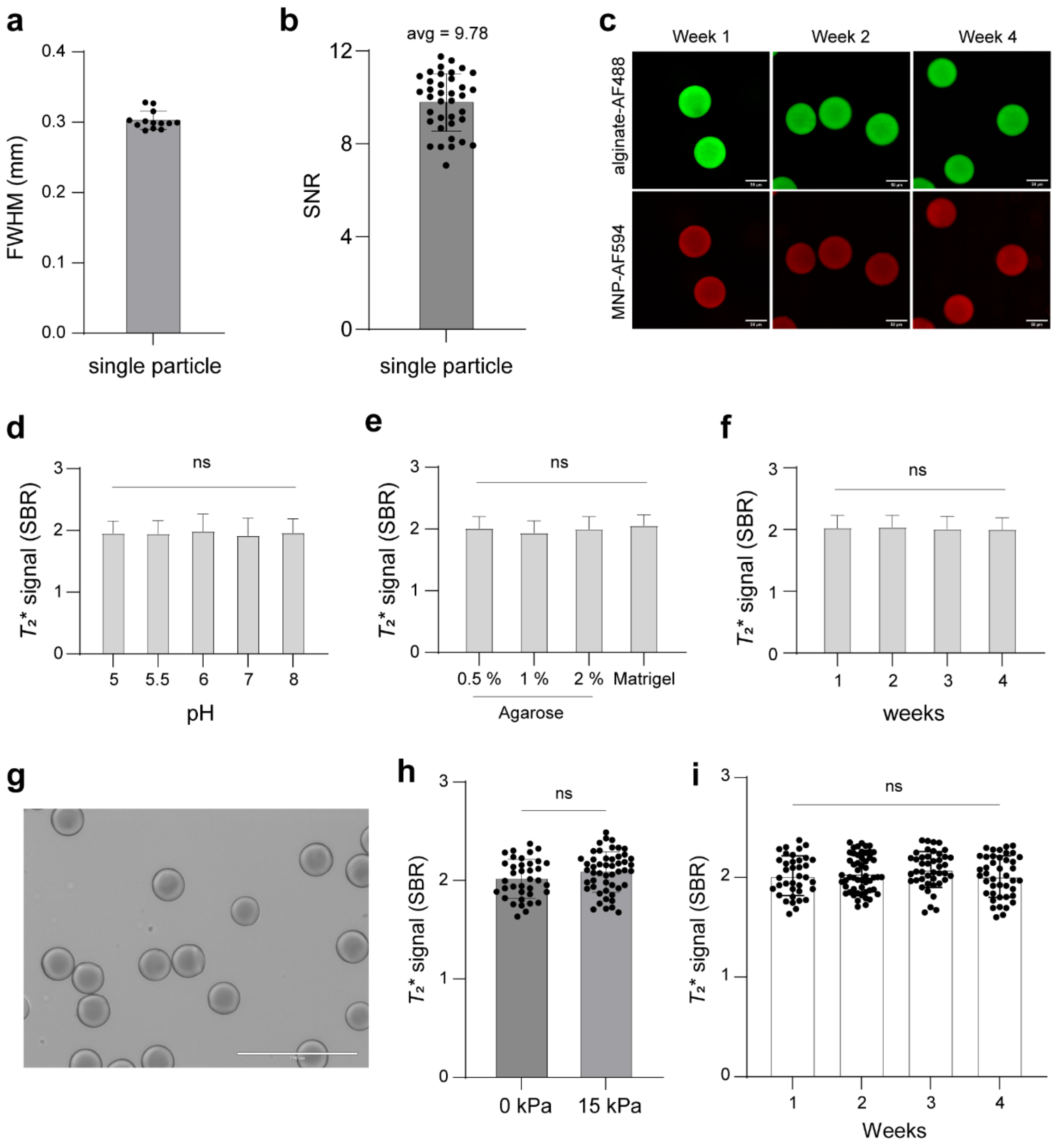


**Extended Data Fig. 1 | Single-particle MRI performance and stability assessment of M$^3$ sensors.**

**a**, Full width at half maximum (FWHM) analysis of single-particle MR intensity profiles used to assess spatial resolution of individual M$^3$ sensors. **b,** Signal-to-noise ratio (SNR) measurements of single-particle MR images, quantifying detection sensitivity under identical acquisition conditions. **c,** Confocal images of M$^3$ sensors at weeks 1, 2, and 4 showing no detectable leakage of MNPs (green, alginate; red, MNPs). **d-f**, $T_2$* signal of M$^3$ sensor measured under different pH conditions (**d**), external matrices (**e**), and over long-term incubation (**f**) for stability testing. **g**, Optical image of PEGDA microparticles. **h**, $T_2$* signal quantification of PEGDA microparticles under 0 and 15 kPa compression, showing no measurable changes in MR signal with applied stress (multiple unpaired two-sided t-test; ns, non-significant). **i**, $T_2$* signal stability of PEGDA microparticles over a 4-week period (one-way ANOVA with Tukey's multiple comparison test; ns, non-significant).

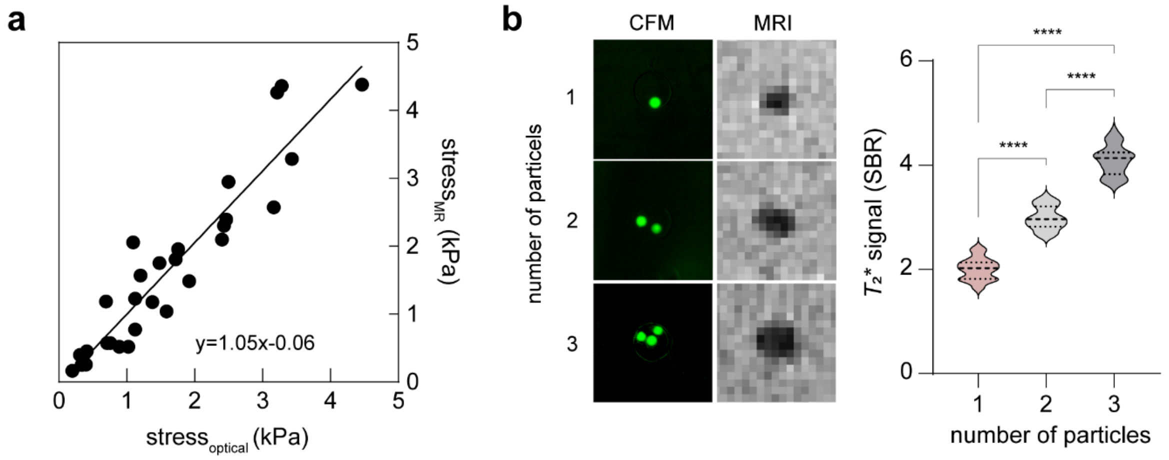


**Extended Data Fig. 2 | Validation of reliable stress measurements using M$^3$ sensors.**
**a**, Correlation between CFM-derived and MR-derived stress measurement in tumor spheroids.
**b**, Representative fluorescence and MR images of different numbers of M$^3$ sensors (left) and corresponding $T_2$* signal quantification (right). (one-way ANOVA followed by Tukey's multiple comparison test;****P < 0.0001).

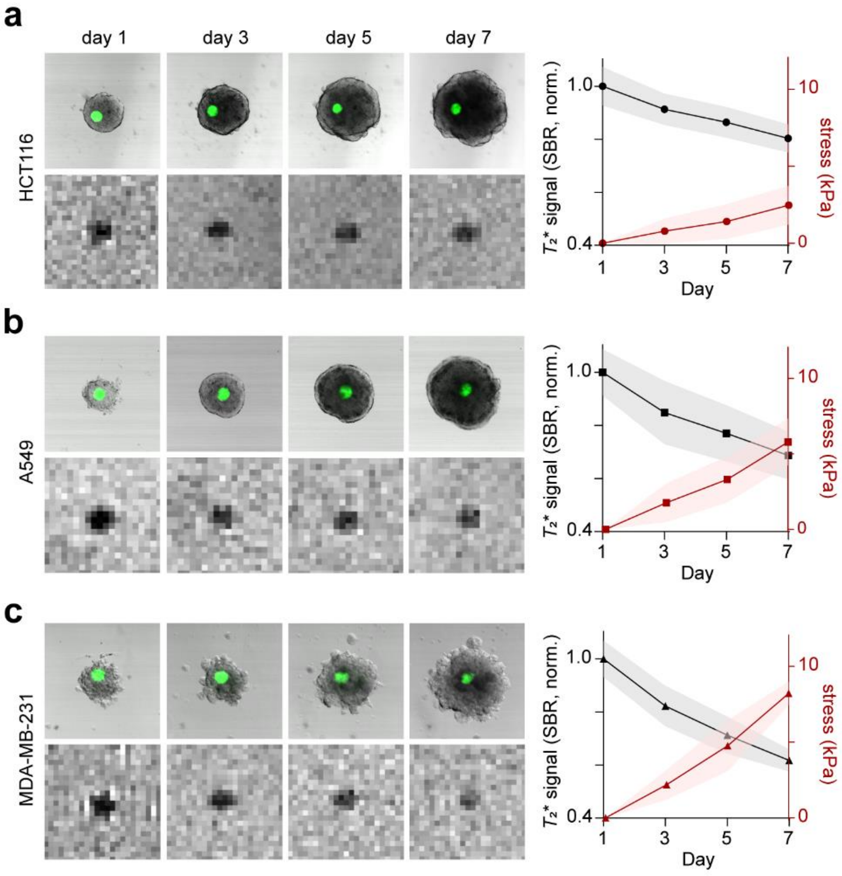


**Extended Data Fig. 3 | Comparison of stress evolution in tumor spheroids from three distinct cancer cell lines.**

**a-c**, Representative CFM and MR image of tumor spheroids formed from (**a**) HCT116 (**b**) A549, and (**c**) MDA-MB-231 (left), and the corresponding normalized $T_2$* signal changes (black) and stress profiles (red) (right).

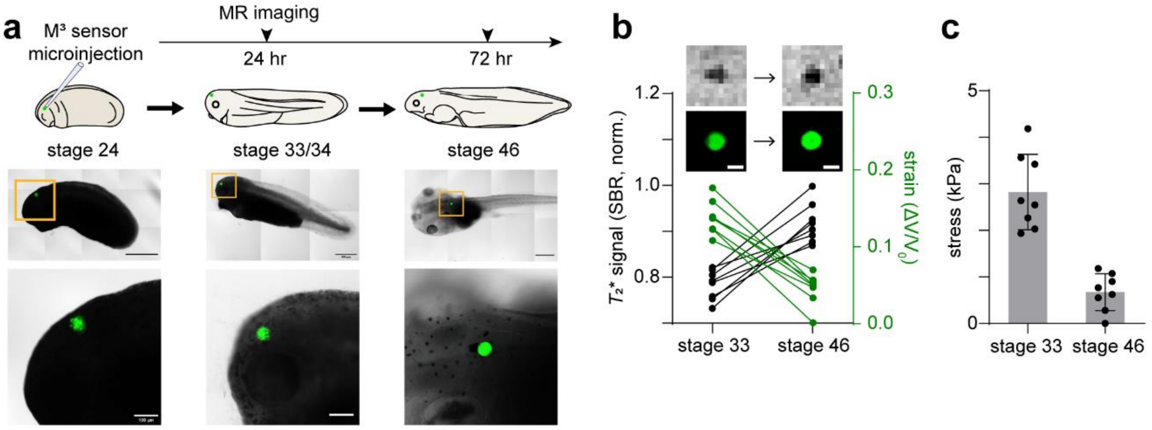


**Extended Data Fig. 4 | Validation of M$^3$ sensor functionality in Xenopus tadpole**.
**a**, Experimental scheme (top) and representative optical images of Xenopus embryos at stage 24 and tadpoles at stage 33/34 and stage 46 (bottom). **b**, Representative CFM and MR images of M$^3$ sensor at stage 33 and stage 46. Normalized $T_2$* signal profiles (left, black) and corresponding strain profiles (right, green) of M$^3$ sensors. **c**, Stress values converted from the MR signals measure by M$^3$ sensors a stage 33 and 46.

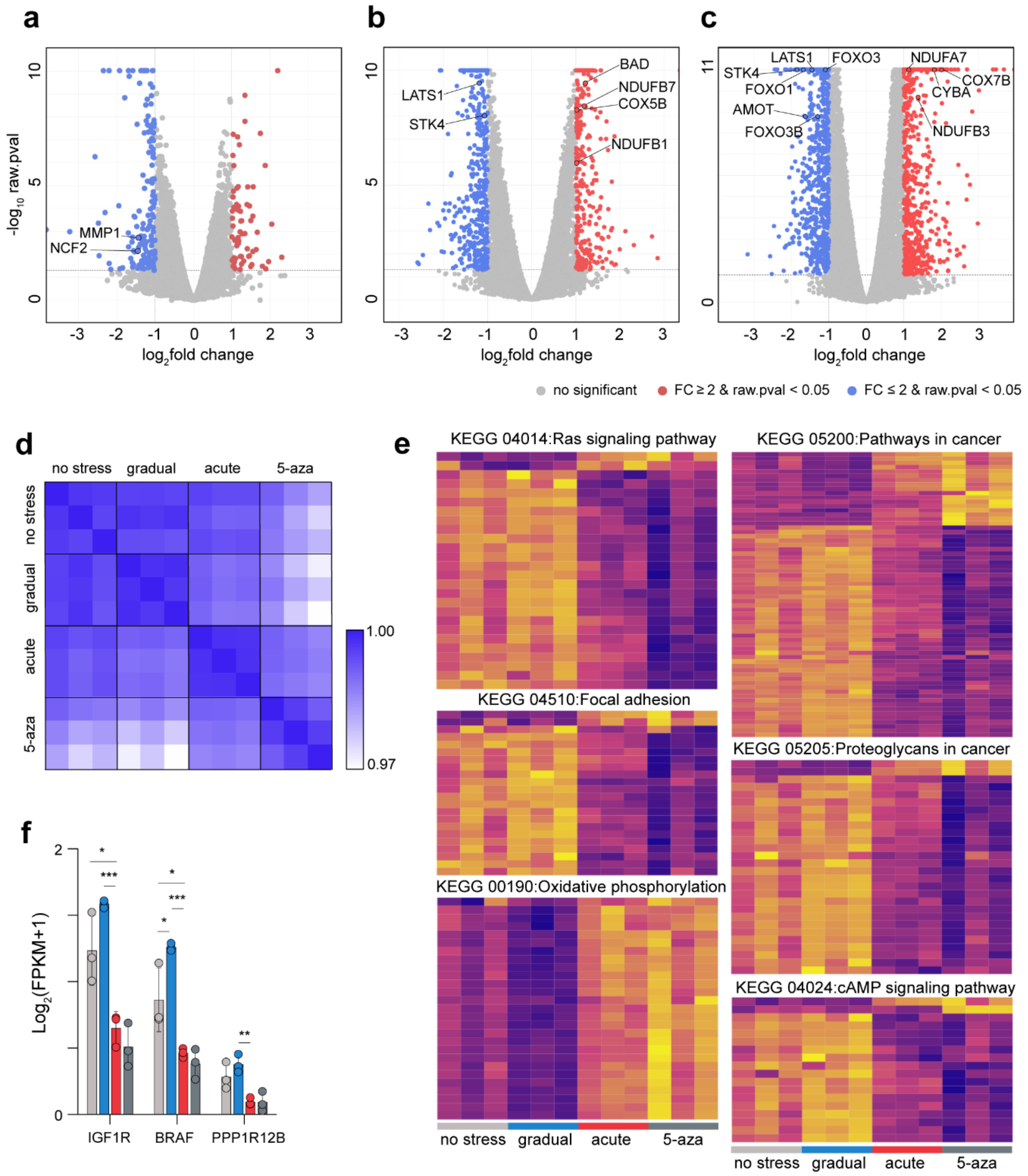


**Extended Data Fig. 5 | Transcriptomic signatures of mechanical stress and epigenetic EMT induction.**

**a-c**, Volcano plots showing differential gene expression between (**a**) gradual stress vs no stress, (**b**) acute stress vs no stress, and (**c**) acute stress vs gradual stress. Representative key genes are annotated for visualization, and significantly upregulated or downregulated genes are highlighted according to adjusted p-value and fold-change thresholds (two-sided Wald test, Benjamini–Hochberg correction). **d**, Pearson's correlation matrix of global transcriptome profiles from no stress, gradual stress, acute stress, and 5-azacytidine-treated spheroids. Samples cluster according to condition, with acute stress showing a close transcriptional similarity to 5-azacytidine treatment, whereas gradual stress remains more similar to the no stress group. **e**, Heatmaps of enriched KEGG pathways, including, Ras signaling (04014),

Pathways in cancer (05200), Focal adhesion (04510), Proteoglycans in cancer (05205), Oxidative phosphorylation (00190), cAMP signaling (04024), highlighting loading stress-dependent gene expression changes. **f**, Comparison of expression values (Log2[FPKM+1]) of selected key genes, including IGF1R, BRAF (RAS-PI3K signaling) and PPP1R12B (actomyosin related regulation).

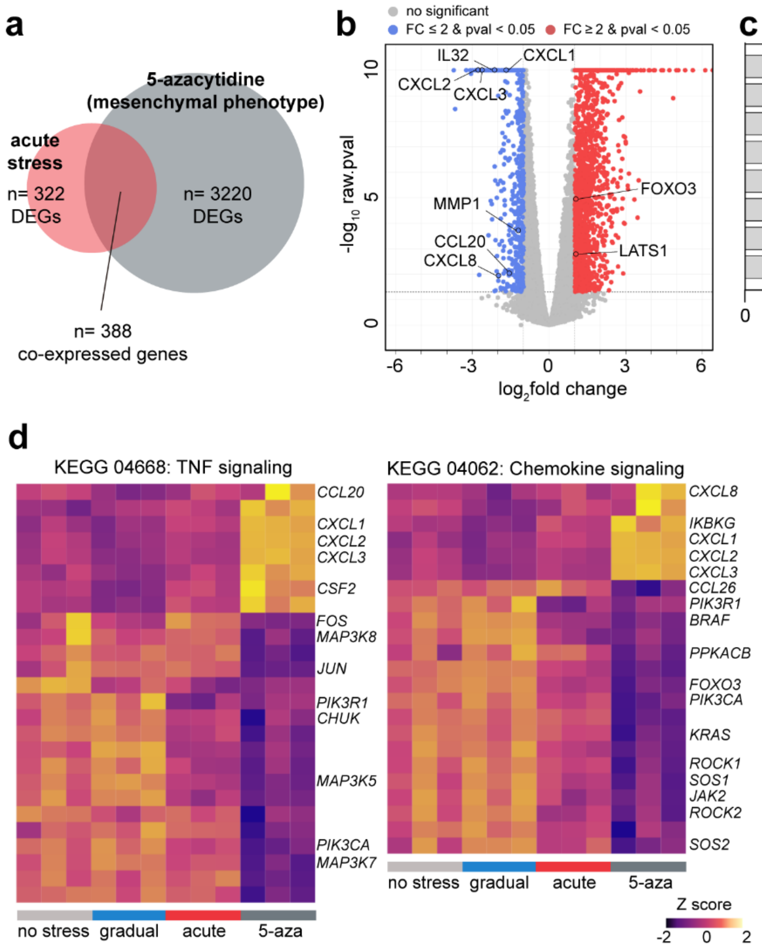


**Extended Data Fig. 6 | Distinct transcriptional signatures reveal limited EMT priming by acute stress compared to epigenetic activation by 5-azacytidine.**

**a**, Euler diagram comparing differentially expressed genes (DEGs) from acute stress vs no stress and 5-azacytidine vs no stress (fold change > 2 and adjusted p < 0.05). A total of 388 DEGs are shared between acute stress and 5-azacytidine, whereas 5-azacytidine uniquely induces 3,220 additional genes, indicating that acute stress triggers only a partial EMT-related transcriptional program (priming), while 5-azacytidine drives a much broader epigenetic EMT shift. **b**, Volcano plots showing differential gene expression between acute stress vs 5-azacytidine. Representative key genes are annotated for visualization. **c**, Bar plot of KEGG pathways specifically enriched in 5-azacytidine–treated spheroids. These pathways reflect inflammation-associated signaling programs, including chemokine signaling, Toll-like receptor signaling, JAK–STAT signaling, NF-κB signaling, cytokine–cytokine receptor interaction, IL-17 signaling, NOD-like receptor signaling, and TNF signaling. **d**, Representative heatmaps of pathways specifically enriched only in 5-azacytidine-treated spheroids, including TNF signaling (04668), Chemokine signaling (04062). **e**, Comparison of expression values (Log2[FPKM+1]) of selected key genes, including IL32, CXCL1, CXCL2, CXCL3, CXCL8, and AREG (inflammatory and chemokine-related genes). These mediators display strong induction exclusively under 5-azacytidine treatment, consistent with epigenetic activation of inflammatory signaling.

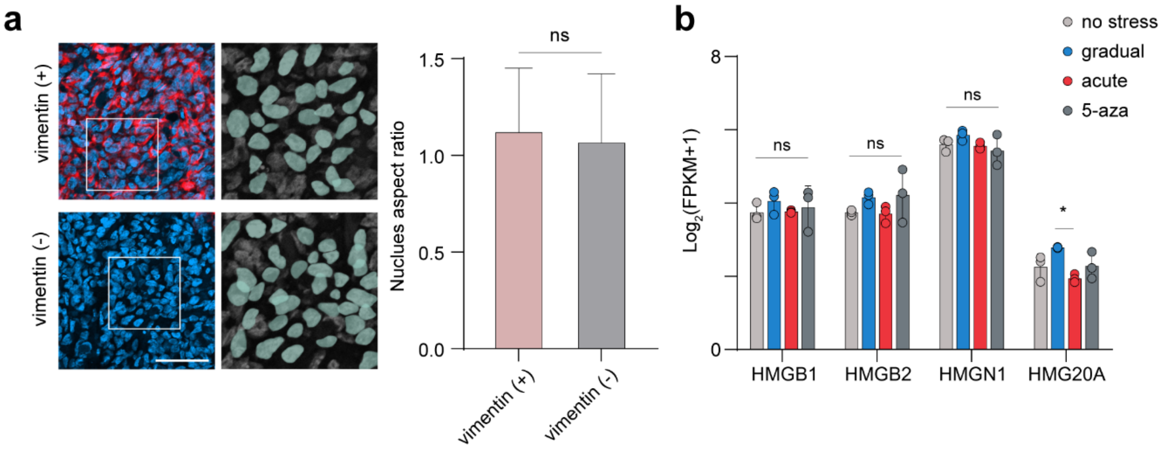


**Extended Data Fig. 7 | EMT induction occurs without nuclear deformation or HMGB2 activation.**

**a**, Representative confocal images of EMT-induced tumor sections showing vimentin-positive and vimentin-negative regions, with magnified views highlighting nuclear morphology (left). Quantification of nuclear aspect ratio (right) shows that nuclei in both regions remain non-elongated. **b**, Comparison of expression values (Log2[FPKM+1]) of HMG family (HMGB1, HMGB2, HMGN1, and HMG20A). No significant changes were observed in the expression of HMGB2 or its related family genes, which are reported nuclear compression–induced EMT drivers.
